# Age-based shifts in multicellular community organization and abundance in HPV-Negative Head and Neck Cancer

**DOI:** 10.64898/2026.10.30.762827

**Authors:** Lina Kroehling, Anthony Spinella, Xaralabos Varelas, Stefano Monti

## Abstract

Aging represents a fundamental driver of cancer risk and progression, yet the mechanisms by which aging reshapes the tumor microenvironment remain incompletely understood. To interrogate how aging alters cellular and microenvironmental organization in cancer, we developed an integrated single-cell atlas of HPV-negative head and neck squamous cell carcinoma (HNSCC) encompassing 73 patients aged 18– 91 years. Using a multi-layered computational framework combining NMF-based module discovery and partial correlation network analysis, we reveal that age-associated differences in tumor composition are driven primarily by selective shifts in cell type abundance and organization rather than widespread age-dependent transcriptional rewiring. We identify an age-enriched cellular community centered on basal-like squamous epithelial states linked to epithelial-mesenchymal transition, matrix-associated myofibroblast-like CAFs, and FOLR2+ macrophages, which secrete immunosuppressive and ECM-remodeling factors. Cross-dataset in-silico validation via TCGA transcriptomes, spatial transcriptomics, and mouse tumor models confirms the correlation of these cell types across patients, specific ligand-receptor usage, and increased proportions of epithelial and stromal cells in aged mouse models. Among epithelial cell-intrinsic age-associated transcriptional changes, midkine (MDK) emerges as a rare but robust tumor-derived signal linked to stromal remodeling, immunosuppression, and invasive epithelial programs. These findings reveal conserved mechanisms of age-dependent cellular reorganization in the tumor microenvironment and identify MDK and associated stromal-remodeling pathways as potential intervention points for age-stratified cancer therapy.

## Introduction

Aging is a risk factor for cancer, yet the mechanisms by which it reshapes the tumor microenvironment remain incompletely understood. Across tissues, aging has been linked to altered immune composition, chronic inflammatory signaling, extracellular matrix remodeling, and impaired tissue repair, all of which may influence tumor initiation, progression, and therapeutic response.^1,2,3,4,5^ However, it remains unclear how aging impacts the cellular organization of human tumors, as well as how these processes synergize with cancer processes to manifest in head and neck squamous cell carcinomas (HNSCC).

Recent single-cell studies have worked to resolve the cellular diversity of HNSCC, identifying malignant epithelial programs linked to differentiation and partial epithelial-to-mesenchymal transition, as well as diverse fibroblast, endothelial, and immune populations.^6,7^ While these studies have provided a framework for understanding intratumoral heterogeneity, age has rarely been treated as a primary axis of variation. More broadly, whether aging acts mainly through shifts in cell-type composition, through rewiring of cell-intrinsic transcriptional programs, or through coordinated remodeling of multicellular communities, and how it affects response to different types of treatments remain unknown. Limiting our understanding is the fact that only 10% of patients in cancer clinical trials are over 75, and most mice used in preclinical studies are 6-8 weeks old, corresponding to a human age of around 20.^1,8^ While immune checkpoint inhibitor (ICI)-based therapy has been shown to be effective in both young and old patients, a recent study showed that despite similar clinical outcomes older patients have altered baseline immune characteristics that behave differently before and during their ICI treatment such as lower cytokine responses and naïve T cell pools.^9^

We recently assembled a single-cell atlas of HPV-negative HNSCC enabling high-resolution characterization of the disease, harmonization of nomenclature, and association of molecular features with clinical phenotypes. Here, we further expanded the atlas to encompass 73 patients aged 18–91 years and implemented a multi-layered analytical framework that combined cell-type identification and age stratification, partial-correlation-based cell-type co-occurrence analysis, and cell–cell communication modeling to interrogate how aging reshapes the tumor TME. This framework enabled systematic identification of age-associated changes in tumor composition, organization, and intercellular interactions. We utilize two external spatial transcriptomic datasets, TCGA and CPTAC data, and mouse models to show our findings can be identified across isolated experiments, data types, and models.

We find that aging is associated less with widespread transcriptional rewiring *within* individual cell types than with selective changes in the abundance and organization of cell subtypes. Older tumors are enriched for a multicellular *community* – defined here as recurrent assemblies of cell-type subsets – centered on basal-like malignant epithelial states, matrix- and myofibroblast-like cancer associated fibroblasts, and macrophages. We further identify midkine (MDK) as a rare but robust age-associated tumor-derived signal linked to these TME-level changes. Together, our results suggest that aging in HPV-negative HNSCC is characterized by coordinated reorganization of tumor–stroma–immune interactions that may lead to poorer outcomes.

## Results

### An Expanded Single Cell Atlas identifies age-associated cellular states with prognostic relevance in HPV-negative HNSCC

We previously established a single cell atlas of HPV-negative HNSCC and here extended this resource by integrating samples from Bill et al.^7^, generating a combined atlas comprising 294,333 cells from 73 patients ranging in age from 18 to 91 years (Figure 1a). To delineate the global cellular landscape, we classified cells into eight major compartments: epithelial and salivary cells, lymphatic endothelial cells, vascular endothelial cells, mesenchymal cells, T and NK cells, B and plasma cells, myeloid cells, and mast cells (Figure 1b). The representation of these compartments varied substantially across patients, consistent with the marked intertumoral heterogeneity that characterizes HPV-negative HNSCC, while also reflecting in part study-specific batch effects. We next performed compartment-specific re-clustering to resolve finer-grained cellular diversity, identifying 89 high-resolution subclusters across the atlas (Figure 1c,d; Supplementary Figures 1 & 2). We performed additional comprehensive characterization of the epithelial compartment, using nomenclature consistent with our previous study^10^. (Supplementary Figure 3).

**Figure 1:**
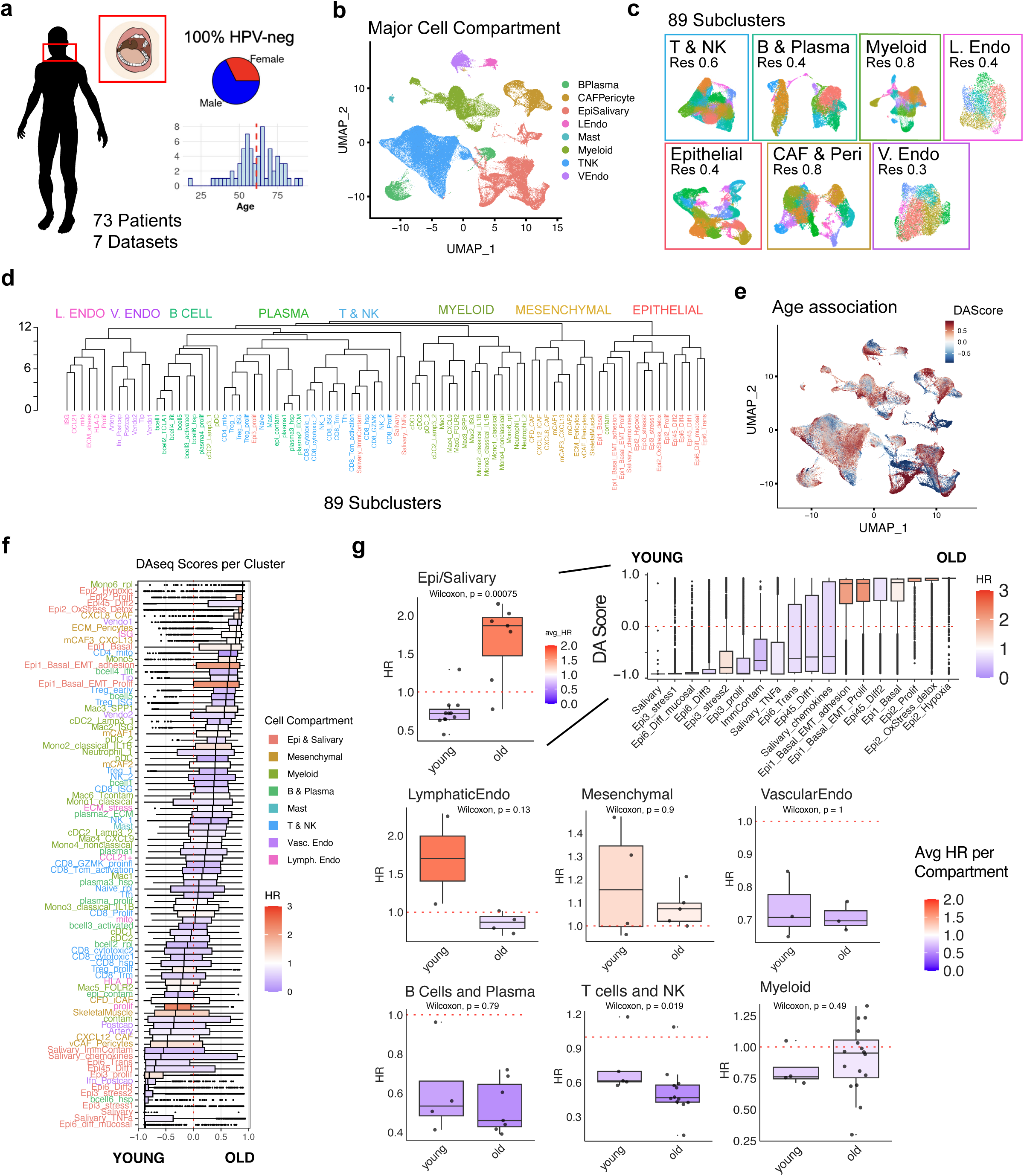
Single cell HPV-negative HNSCC atlas. a) Summary of sex distribution and age range of 73 patients included in the atlas. b) Cells colored by major cell compartment. c) High resolution clustering of each of the major cell compartments for a total of 89 cell sub-types. d) Hierarchical clustering of the 89 cell sub-types, colored by major cell compartment. e) Atlas UMAP colored by DAseq age prediction score. f) Boxplots showing the average DASeq score of each cell per 89 cell sub-types, colored by hazard ratio of the cell sub-type signature in the TCGA dataset. g) Boxplots showing the average HR across the 89 cell sub-type signatures in the TCGA dataset, split by if the average DAseq score of the given cluster is below 0 (young) or above 0 (old). P-values from wilcoxon test.

Next, we examined aging-induced transcriptional changes within each compartment. To improve robustness, we used two complementary differential expression strategies. First, we applied MAST to model gene expression as a function of age at the single cell level, while including patient and dataset as random effects to account for pseudoreplication bias (the non-independence of cells from the same patient). Second, we performed patient-level pseudobulk differential expression within each compartment using edgeR, with dataset and sex included as fixed effects. Across compartments, age-associated differential expression was limited (Supplementary Fig. 4a), consistent with prior studies showing that bulk expression differences primarily reflect altered cellular composition rather than widespread cell-intrinsic rewiring^11^.

Midkine (MDK) emerged as the only gene consistently significantly upregulated in epithelial cells from older patients (Supplementary Figure 4b). MDK has previously been linked to older age in healthy individuals^12^ and in cancer^13^, and has been implicated in aggressive HNSCC and linked to immune evasion^14^. However, these observations have not been linked in the context of age-associated HNSCC biology. MDK was expressed across multiple epithelial subclusters (Supplementary Figure 4c-d) and was also elevated in older HPV-negative HNSCC tumors in TCGA (p = 0.0052, Supplementary Figure 4e). Notably, Pleiotrophin (PTN), another secreted heparin-binding growth factor with approximately 50% sequence similarity to MDK, was also upregulated in the older epithelium, although this association was detected only in the pseudobulk analysis.

We then looked for age-associated shifts in cell type proportions in a cell-type agnostic manner. Using DA-seq, which does not depend on pre-identified clusters, we identified widespread age-enriched cellular neighborhoods across multiple compartments (Figure 1e-f), indicating that aging is associated with the selective expansion or shrinking of specific cellular subtypes or states rather than uniform changes in major lineage abundance.

To assess the clinical relevance of these subtypes, we projected the 89 high-resolution cell subtype signatures onto bulk RNA-seq profiles from 420 HPV-negative HNSCC tumors in TCGA^15^, and tested each signature for association with overall survival using Cox proportional hazards models. Because no signatures reached fdr significant associations with HR, we looked at HR trends within age groups in each compartment. Signatures derived from B cells, plasma cells, and CD4/CD8 T-cells were generally associated with lower hazard ratios, whereas signatures from malignant epithelial and proliferative states were associated with higher hazard ratios (Figure 1f), consistent with a protective role for adaptive immune programs and an adverse prognostic effect of aggressive tumor-intrinsic states. Among all compartments, epithelial and salivary cells showed the largest difference in hazard ratios between young- and old-enriched states (pval = 0.00075), while other groups showed more modest trends in the same direction (mesenchymal HR >1 in both young and old, and B cells HR < 1 in both young and old) (Figure 1g). Within the epithelial compartment, the subpopulations most strongly associated with poor survival comprised squamous lineage basal-like states enriched for EMT and adhesion programs (Epi1_basal_EMT_adhesion & Epi1_basal_EMT_prolif), as well as another squamous lineage basal-like cluster, Epi2, a state characterized by drug metabolism and oxidative stress signatures (Supplementary Figure 3e). Interestingly, Epi1_basal_EMT_adhesion, Epi2_OxStress_Detox, Epi2_Hypoxia, and Epi2_prolif all contained a small portion of cells with high expression of LGR5, suggesting these could harbor small populations of more plastic tumor cells. The Epi3 clusters (Epi3_stress1, Epi3_stress2 and Epi3_prolif), enriched for TNFa signaling via NFKB and the AP1 pathway, were enriched in younger patients, along with salivary cells and three out of four of the more differentiated squamous cell clusters. These findings suggest that aging may influence prognosis in part by shifting the balance between protective immune states and high-risk tumor cell programs.

### Integrative network analysis reveals age-enriched cellular communities

Rather than focusing on changes in individual sub-populations, we decided to look for groups of sub-populations changing in proportion together across patients, which would suggest coordinated shifts in multicellular communities within the tumor microenvironment that we could then associate with age. To identify such communities, we developed an integrative approach that combines two complementary strategies to detect cell subtypes that co-vary across patients. First, we identified communities of co-proportional cell types using NMF-based module discovery through CoVarNet^16^.

Because the NMF-based module discovery will identify both direct and indirect interactions, we used a dataset-adjusted partial-correlation network built with RSNet^17^ to inferred only *direct* interactions within these communities, where nodes represent cell subtypes and edges represent partial correlations between cell subtype proportions, corrected for batch effects (Fig. 2a–d; Methods). Lastly, we incorporated age information from DAseq and overlaid spatial data to identify age-associated communities with independent external supporting evidence for spatial colocalization. Using this approach on the 89 atlas cell subtypes, we identified 20 communities (CMs) of cell subtypes that both co-vary in proportion across tumors and are directly connected in network space, forming coordinated cellular neighborhoods poised for interaction in the microenvironment (Figure 2b).

**Figure 2:**
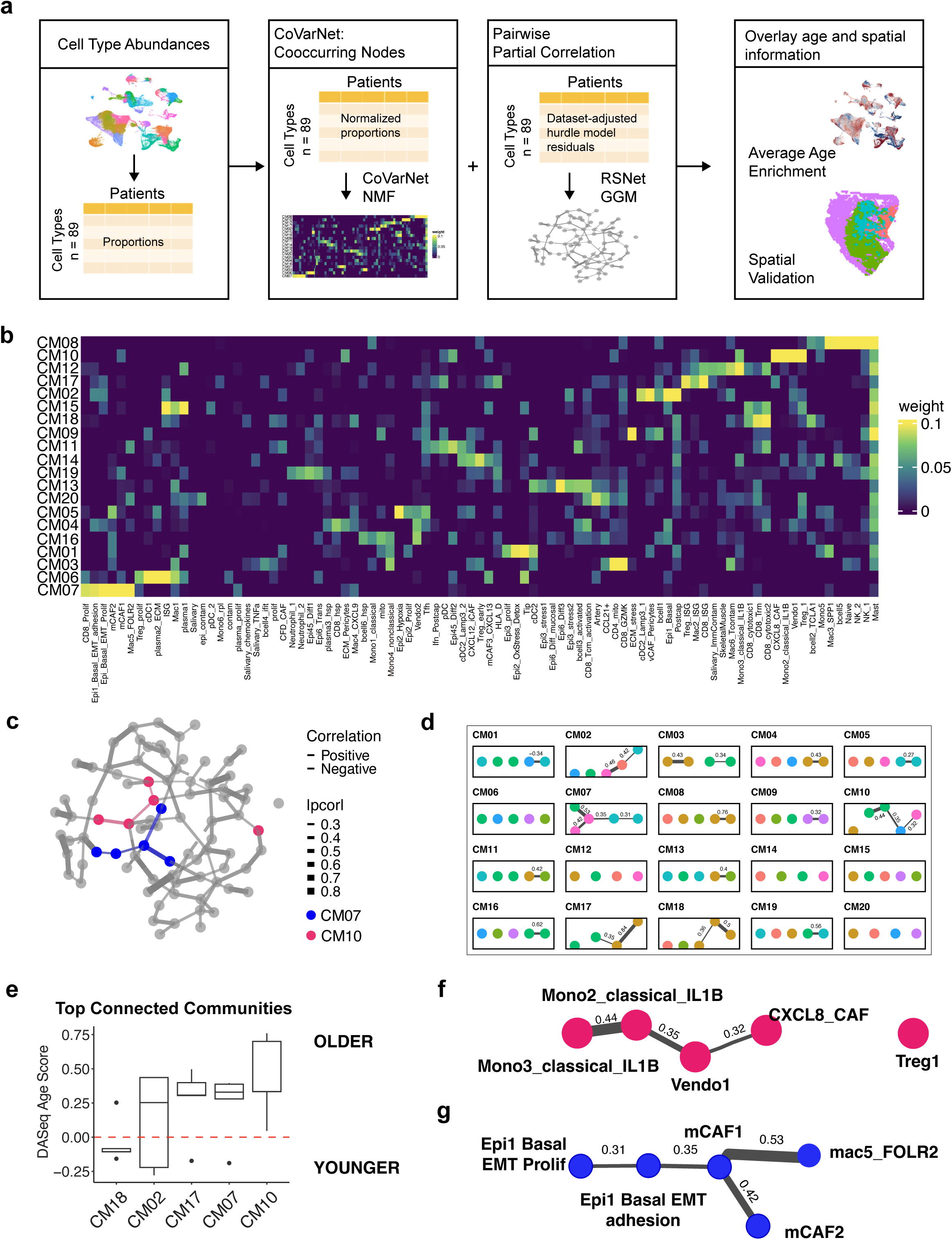
Identification of cell type communities. a) Schematic showing application of NMF via the CoVarNet package and partial correlation network method to detect populations of cells changing in proportion together across patients. b) Heatmap showing weights for each cell sub-type contributing to each community. c) Full partial correlation network, where each node is a cell sub-type and edges shown are partial correlation weights with fdr < 0.05. c) Sub-graph from partial correlation network showing significant edges between top five nodes of each CM. Nodes (cell sub-types) are colored by major cell type classification. e) Boxplots of DAseq scores across top five cell sub-types in each of the communities with at least two edges in the partial correlation network. f) Top five cell sub-types contributing to CM10 and their connectivity in the partial correlation network. Edges shown have fdr < 0.05. g) Top five cell sub-types contributing to CM07 and their connectivity in the partial correlation network. Edges shown have fdr < 0.05.

From each community, we extracted the five top-contributing cell subtypes with the highest NMF weights, and extracted their edges from the partial correlation network (Fig. 2c-d). Five communities contained at least two edges among their top contributors (Fig 2d), which we then focused on, as those with edges are more likely to represent biologically coherent communities with direct communication rather than indirect co-fluctuation driven by shared correlation with a third factor. Among these, four highly connected communities – CM02, CM07, CM10, and CM17 – showed elevated DA-seq age-enrichment scores and were therefore enriched in older patients, whereas CM19 was enriched in younger patients (Fig. 2e). The two most aged-enriched communities showed distinct cellular composition: CM10 comprised of two IL1B+ monocyte clusters, a vascular endothelial cluster, CXCL8+ CAFs, and Tregs, whereas CM07 consisted of the two malignant epithelial clusters previously described (Epi1_Basal_EMT and Epi1_Basal_EMT_Prolif), two mCAF clusters (mCAF1 and mCAF2), and a macrophage cluster marked by FOLR2. (Fig 2f-g).

### An age-linked tumor–mCAF–macrophage community associated with immunosuppression and stromal remodeling

Among the age-associated communities, CM07 was of particular interest because it was enriched in older patients, exhibited the highest internal connectivity, and contained epithelial sub-populations with HR’s trending towards worse survival as we previously showed.

The myeloid cluster Mac5_FOLR2 had the highest weight contributing to the community (Figure 3a). Notably, the UMAP (Supplementary Figure 2b) and hierarchical clustering (Figure 1d) showed that Mac5_FOLR2 is closely related to SPP1^+^ and CXCL9^+^ macrophage subpopulations, which have recently been described as macrophage polarization states with prognostic relevance in HNSCC.^7^ Although Mac5_FOLR2 shared expression of genes such as TGFBI, MMPs, and APOE with these related populations, it was distinguished by higher expression of FOLR2, CXCL1, CXCL2, and CXCL3, suggesting functional divergence despite partial transcriptional overlap (Figure 3b). FOLR2 is a marker of tissue residency, and its expression has been identified in tissue-remodeling macrophage states described in other cancer and aging tissues, including FOLR2^+^ macrophages implicated in fibrotic communities^18^ and tumor immunosuppression.^19^ Although not significantly differentially expressed in the atlas, FOLR2 was upregulated in older patients in the HPV-negative HNSCC TCGA subset of patients (p-value = 0.0013) (Figure 3c). Pathway overrepresentation analysis highlighted not only canonical macrophage programs, such as complement cascades (also enriched in the CXCL9 macrophage population), but also chemokine and cytokine signaling, which was unique to the Mac5_FOLR2 population (Figure 3d). In addition, Mac5_FOLR2 was enriched for focal adhesion and ECM receptor interactions (as was the SPP1+ population), and regulation of actin cytoskeleton pathways, consistent with adaptation to matrix-rich stromal environments (Figure 3d).

**Figure 3:**
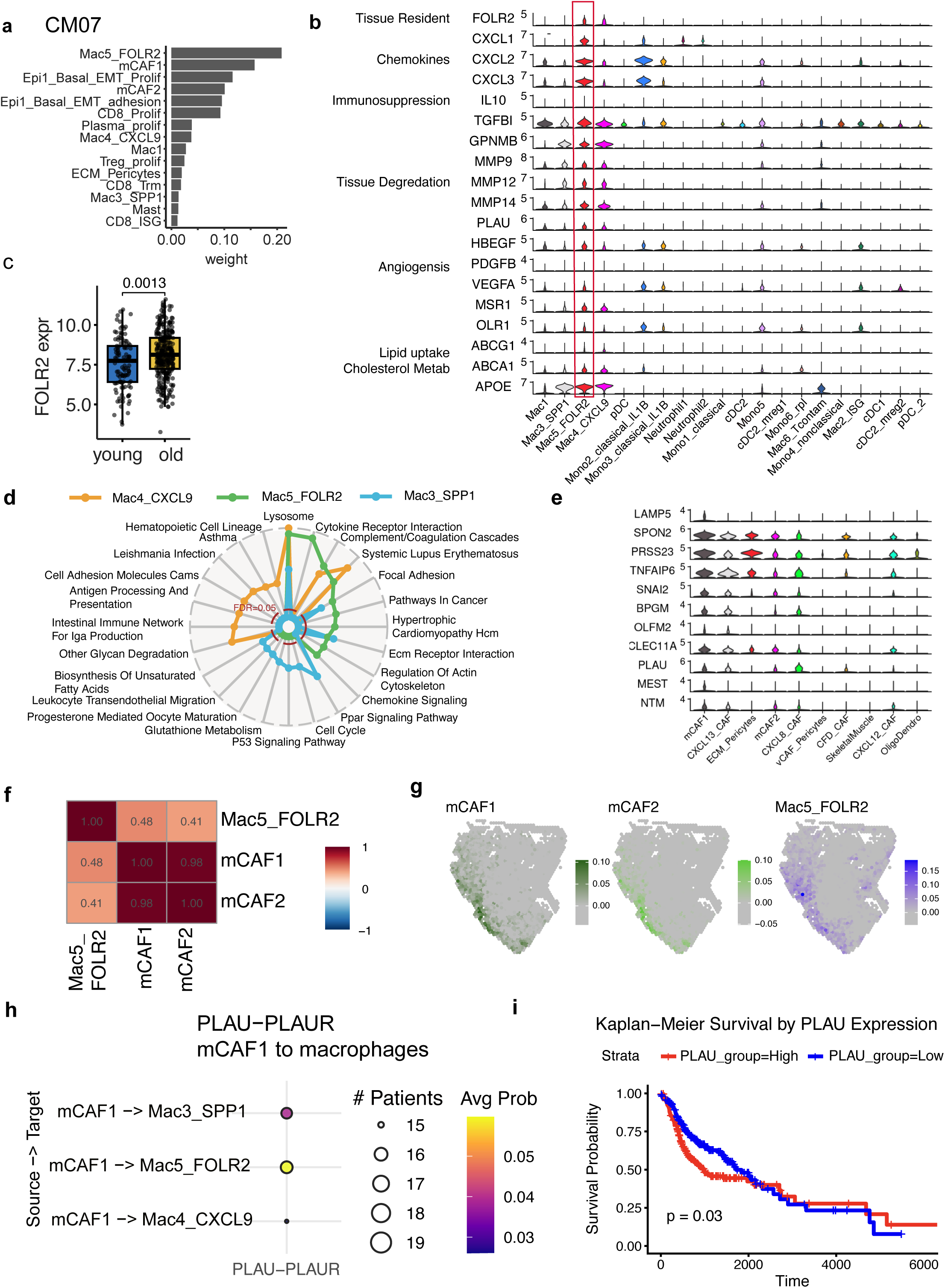
An epi-CAF-myeloid age-linked signaling hub. a) Barplot showing top 15 cell sub-types contributing to CM07 and their weights. b) Violin plot showing marker genes for Mac5_FOLR2 population in red with respect to all other myeloid populations. c) Boxplot showing FOLR2 expression in HPV-negative HNSCC TCGA samples. Young < 65 and Old >= 65; Wilcoxon p-value = 0.0013. d) Spider plot showing top 10 enrichment terms for Mac4_CXCL9, Mac3_SPP1 and Mac5_FOLR2 populations. The red dotted line shows the fdr < 0.05 cutoff. e) Stacked volcano plot showing marker genes for mCAF1 population with respect to all other mesenchymal populations. f) Correlation of mCAF1, mCAF2, and Mac5_FOLR2 signatures in Arora spatial data across patients. g) Representative spatial sample showing projection of atlas mCAF1, mCAF2, and Mac5_FOLR2 signatures. h) Specific PLAU pathway ligand receptor interactions being sent from mCAF1 to macrophage clusters. Bubble size represents the number of patients the interaction was identified in, and the color corresponds to the average probability of the interaction across patients. i) Kaplan-Meier survival analysis by PLAU expression in HPV-negative HNSCC TCGA dataset.

Within the partial correlation network, Mac5_FOLR2 was directly connected to mCAF1 (partial correlation = 0.53, FDR = 7.11e-8), the population contributing the second highest weight to the community, which in turn was connected to mCAF2 (Figure 2g). Relative to other fibroblast and pericyte subclusters, mCAF2 was enriched for extracellular matrix (ECM) receptor interaction, focal adhesion pathways, and a myofibroblast-like CAF signature (Supplementary Figure 5a,b), whereas the mCAF1 population was distinguished by expression of LAMP5 and LGALS9 (Figure 3e), an ECM matrix-like CAF signature, and by enrichment for cycling CAFs, p53 signaling, and cytokine-cytokine receptor interaction pathways (Supplementary Figure 5a-b). These features are consistent with fibroblast populations described in other tumor types, including a TGFβ-associated myofibroblast-like population in breast cancer associated with an immunosuppressive microenvironment.^20^ Although mCAF1 and mCAF2 were not the most age-enriched mesenchymal cell sub-types, both showed a bias toward older patients and occupied central positions in the network.

We utilized an independent spatial dataset to infer proximity of the mCAF1 and Mac5_FOLR2 populations. Across 12 spatial tissue samples, pairwise correlation analysis revealed a positive association between the Mac5_FOLR2 and mCAF1 (estimate = 0.525, Pr(>|t| < 2^-16^), with a slightly stronger correlation to mCAF1 than mCAF2 (Figure 3f, Supplementary Figure 5c). Using a linear model associating the signatures while including a random effect term for sample confirms these positive associations. In a representative tissue section, both CAF and macrophage populations colocalize around the tumor edge (Figure 3g). Additionally, the marker genes for these populations were also correlated in the spatial data (Mac5_FOLR2 signature and FOLR2 gene; estimate = 1.5, fdr = 6.14^-107^; mCAF1 signature and LAMP5 gene estimate = 1.16, fdr = 5.94^−96^) (Supplementary Figure 5d).

We then asked if these populations were connected through specific signaling pathways. Relative to other macrophage populations, mCAF1-to-Mac5_FOLR2 communication was characterized by strong collagen, APP, MIF, PLAU, GALECTIN and MHC-I signaling (Supplementary Figure 5e). The PLAU-PLAUR interaction was particularly strong relative to analogous fibroblast-to-macrophage interactions, as measured by interaction probability (Figure 3h). The spatial data showed higher correlation of PLAUR with the Mac5_FOLR2 signature than with the SPP1+ and CXCL9+ signatures (Supplementary Figure 5d), suggesting that this macrophage state may be especially poised to receive fibroblast-derived suppressive or matrix-associated signals. We looked for external validation for correlation of the marker genes of these cell types, and see significant correlations between FOLR2 and PLAUR and the Mac5_FOLR2 population, and PLAU and LAMP5 with the mCAF1 population in the HNSCC CPTAC dataset (Supplementary Figure 5f). Notably, PLAU is associated with worse survival in the TCGA dataset (Figure 3i).

The other two cell types contributing most to CM07 and connected in the partial correlation network were Epi1_Basal_EMT_adhesion and Epi1_Basal_EMT_Prolif. These basal-like squamous epithelial subclusters were enriched for the tumor leading-edge signature described by Arora *et al.*^22^, whereas differentiated epithelial subclusters assigned to other communities, were enriched for tumor-core programs (Fig. 4a-b). This pattern is consistent with the model proposed by Puram *et al.*^6^, in which partial-EMT tumor cells localize to the invasive front and interact with CAFs, whereas more differentiated tumor cells occupy the tumor core. Supporting these spatial relationships, the mCAF1 signature positively correlated with the Epi1_Basal_EMT_adhesion signature in the Tan tumor and TME datasets (Figure 4c).

**Figure 4:**
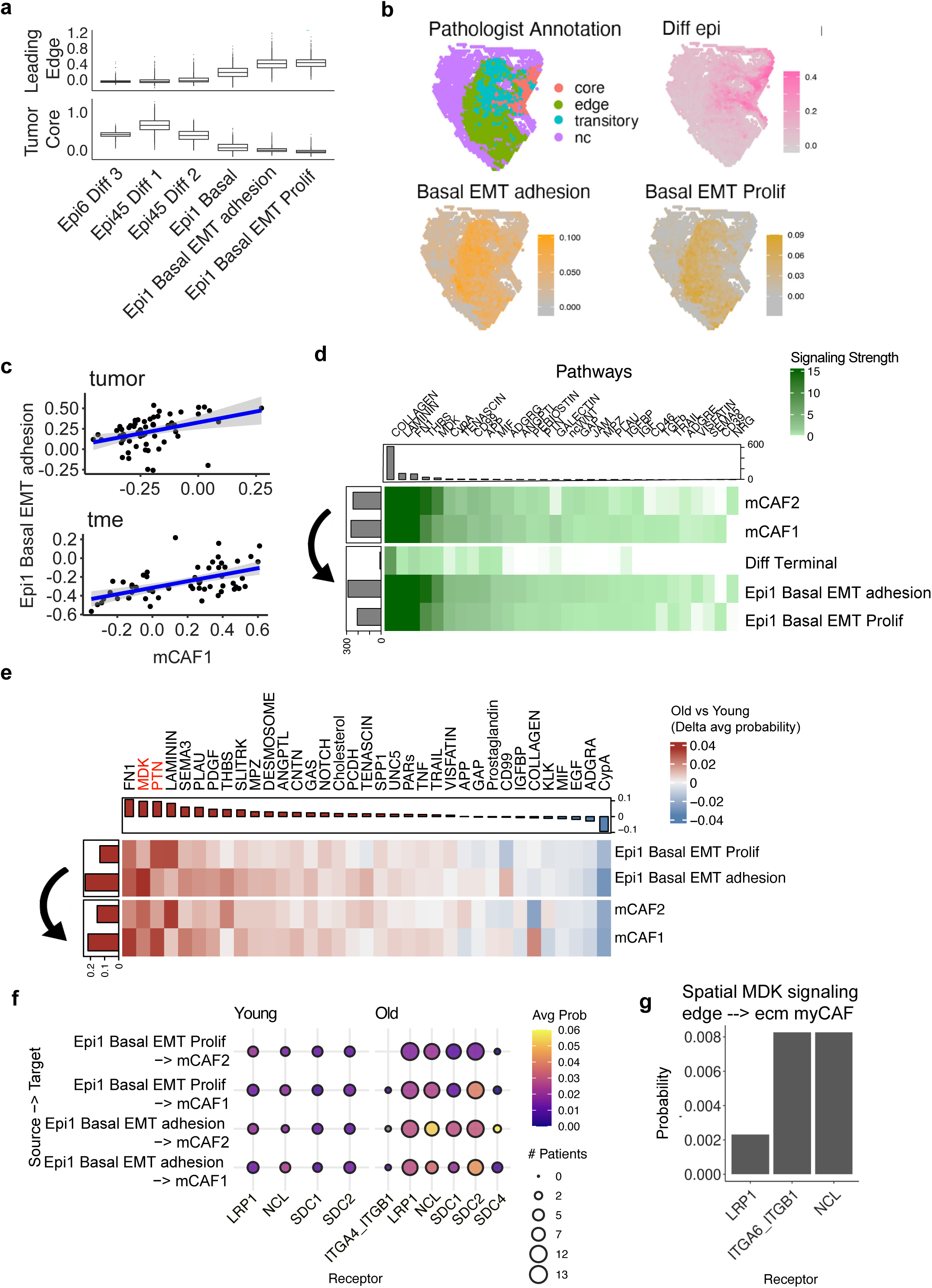
An epi-CAF-myeloid age-linked signaling hub. a) Boxplots showing expression of tumor core and leading edge signatures on epithelial and salivary cell sub-types. b) Representative spatial sample showing projection of atlas epithelial CM07 signatures. c) Correlation of Epi1_Basal_EMT_adhesion and mCAF1 signatures in tumor (top) and TME (bottom) samples from Tan et al. d) Summary of signaling pathways being sent from mCAF1 and mCAF2 to basal and differentiating epithelial cell sub-types to. Color shows predicted interaction strength for the given pathway. e) Differential heatmap showing difference in signaling pathways between old and young patients in signals being sent from epithelial cells to mCAFs. Red indicates pathway signaling probability is higher in older patients, blue indicates signaling is higher in younger patients. f) Specific MDK ligand receptor interactions being sent from epithelial clusters to mCAF1 and mCAF2. Bubble size represents the number of patients the interaction was identified in, and the color corresponds to the average probability of the interaction across patients. g) Signaling probability being sent from tumor edge to ecm.myCAFs from MDK to different receptors. (Data from Arora et al.)

We next evaluated whether the inferred co-occurrence of CM07 populations reflected intercellular communication. To reduce false-positive interactions from cross-patient pooling, we performed CellChat separately for each patient at the 89-subtype resolution and aggregated the results across patients. Comparison of mCAF1 and mCAF2 signaling toward basal-like vs. differentiated epithelial subclusters revealed markedly stronger signaling to the basal-like cells across multiple pathways, consistent with their leading edge signature enrichment (Figure 4d). These interactions were dominated by ECM- and adhesion-related pathways, including laminin, collagen, and fibronectin signaling.

Because both epithelial and mesenchymal populations in CM07 were enriched in older age, we asked whether specific signaling pathways showed elevated activity in older patients. Many pathways displayed greater signaling in older vs. younger patients, with FN1, MDK, PTN, and LAMANIN among the most increased (Figure 4e).

Examining ligand-receptor pairs within the MDK pathway revealed significantly elevated signaling probability between multiple pairs in older patients, suggesting MDK as a potential mechanism linking age-associated tumor states to stromal organization. (Figure 4f). Cell cell communication analysis in the independent spatial dataset also identified MDK-ITGA6/ITGB1 and MDK-LRP1/NCL signaling between tumor edge cells and ecm.myCAFs, a population identified in Arora et al., which overlaps with our myCAF1 and myCAF2 signatures. (Figure 4g), although the spatial dataset’s modest size limits robust age stratification.

Collectively, these findings suggest that invasive epithelial states, activated fibroblasts, and macrophages expand in concert across tumors, acting as a coordinated age-linked multicellular community.

### Age-associated programs are partially conserved in mouse oral carcinogenesis models

We previously explored age-associated changes using the 4MOSC1/4NQO mouse model of oral carcinogenesis. We reanalyzed this data to validate age-associated cell subtype proportion changes observed in the human atlas. In the atlas, the CXCL12_iCAF population, most consistent with a canonical iCAF state, was enriched in younger patients, whereas mCAF states were more enriched in older patients (Figure 5a). This pattern mirrored the mouse mesenchymal compartment, in which young tumors showed high iCAF program activity, whereas older tumors displayed higher mCAF program expression (Figure 5b). Although CXCL8+ CAFs were also enriched in older human tumors, this population diverges from canonical iCAF programs, as we have previously shown.^10^

**Figure 5:**
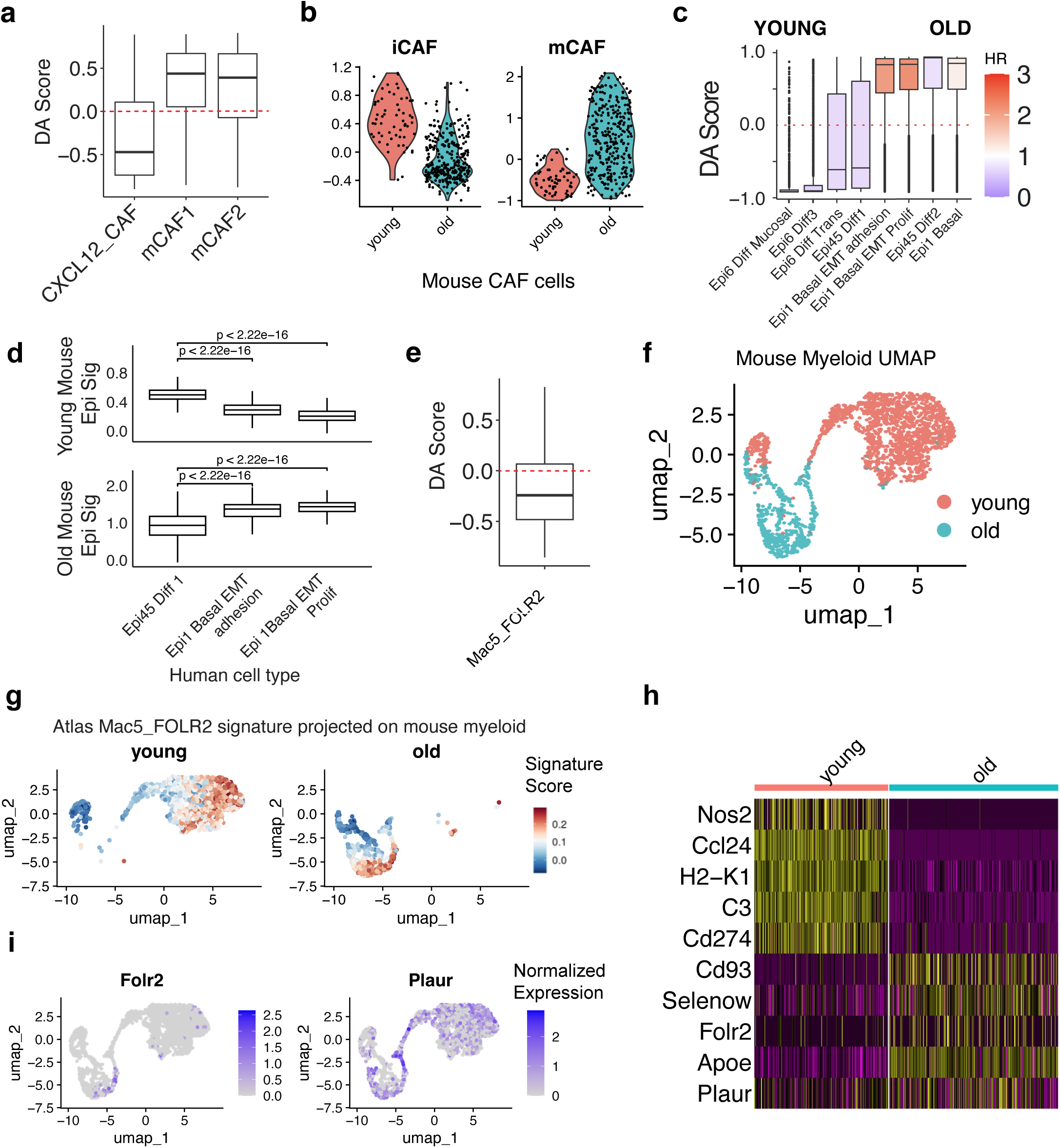
Mouse data. a) Boxplots showing DAscore of iCAF and myCAF populations in atlas. b) Violin plots showing iCAF and myCAF signature expression in fibroblasts in young and old mice. c) Boxplots showing DAseq scores for the basal and differentiated epithelial cell sub-types. d) Boxplots showing the old mouse epithelial signature (top) and young mouse epithelial signature (bottom) projected across the differentiated, Epi1 Basal EMT adhesion, and Epi1 Basal EMT proliferating cell sub-type signatures of the atlas. e) Boxplot showing DAscore for the Mac5_FOLR2 cluster. f) UMAP of the mouse myeloid compartment split by age. g) UMAP of the mouse myeloid compartment, colored by expression of the Mac5_FOLR2 signature. h) Heatmap showing differential genes between the old myeloid and young myeloid cells expressing the Mac5_FOLR2 signature. i) UMAP with expression of Folr2 (left) and Plaur (right).

Within the human epithelial compartment, basal-like cell type sub-populations enriched in older patients contrasted with differentiating epithelial cells more prevalent in younger patients (Figure 5c). Projection of the atlas onto age-stratified mouse epithelial signatures recapitulated this pattern, with the young mouse signature mapping to differentiating atlas cells and the older mouse signature to basal-like cells (Figure 5d).

In the human atlas, the Mac5_FOLR2 subcluster showed a trend toward enrichment in younger patients (Figure 5e). Projection of the human signature onto the mouse data identified distinct young and old mouse myeloid populations with strong similarity (Figure 5f-g). Age-dependent transcriptional differences emerged: older mice showed higher expression of Cx3cr1, Cd83, Cd93, Cd72, and Selenop, whereas younger mice expressed higher levels of Cd274, Ccl24, C3, H2-K1, and Nos2 (Figure 5h). Notably, Folr2 was upregulated in the old myeloid compartment, which also highly expressed Plaur, mirroring the human atlas (Figure 5i). Although a Mac5_FOLR2+ related macrophage population was present in both age groups, its functional state appeared to shift with age.

In summary, independent validation in a mouse model confirms that advancing age is associated with coordinated shifts in cell state composition across epithelial, mesenchymal, and immune compartments.

## DISCUSSION

In this study, we integrated single-cell, bulk and spatial transcriptomics, and mouse studies to examine the effects of aging in HPV-negative HNSCC. We found that age-associated differences were driven primarily by shifts in the abundance and co-occurrence of specific cell states rather than widespread transcriptional rewiring within individual states. Specifically, older tumors were enriched for basal-like squamous epithelial programs linked to the tumor leading edge, EMT, and adhesion, which are associated with poor prognosis.

One contribution of this work is the development of a computational framework to identify age-associated communities. The use of partial correlation networks offers a key advantage: distinguishing direct associations between cell subtypes by accounting for indirect correlations driven by shared factors, while utilizing cell type proportion residuals regressed on dataset-specific covariates controls for batch effects, enabling robust cross-dataset comparisons. Additionally, leveraging multiple independent spatial datasets, TCGA, and a mouse model provides robust support for validation of the population’s colocalization and age-biased composition.

Using this framework, we identified an age-associated community of tumor cells, myofibroblast-like and matrix associated CAFs, and macrophages that expands on established CAF-tumor leading-edge biology. In HNSCC, LGALS9+ CAF populations have previously been implicated in promoting CD8 T-cell exhaustion.^23^ Our data further suggest that the role of these mCAFs may extend beyond lymphocyte dysfunction to organization of a myeloid-rich stromal environment, as indicated by the strong network connection between mCAF1 and Mac5_FOLR2. The FOLR2+ macrophage population extends the recently described SPP1/CXCL9 macrophage polarization axis and implicates FOLR2+ macrophages in age-associated stromal remodeling in HNSCC. FOLR2+ macrophages have been localized to inflammatory tumor stroma in xenograft mouse models^24^ and exhibit immunosuppressive functions in other tumors.^25^ Interestingly, FOLR2^+^ macrophages have also been identified as senescent macrophages enriched in old mice with lung tumors, where their ablation decreases tumor burden by promoting immune regulation and reducing vasculature.^26^ We show that FOLR2 is highly specific to the macrophage compartment, which may provide a therapeutic target for folate-conjugated therapies.^25^ Whether and how this population changes with age warrants further investigation.

Beyond identifying signaling pathways within this community, we demonstrate its age association, which is partially recapitulated in mouse models. Matrix-associated myofibroblast-like CAF states were relatively enriched in older tumors, while younger tumors retained more inflammatory-like fibroblast programs. This shift is biologically plausible given established links between aging, altered wound healing^27,28^, and extracellular matrix deposition.

Among the relatively few cell-intrinsic transcriptional differences associated with age, MDK emerged as a candidate tumor-derived signaling node connecting stromal remodeling and immune suppression. MDK has been linked to cisplatin resistance in HNSCC cell lines^29^, suggesting that cisplatin treatment could be paired with MDK inhibitors to improve treatment response. MDK inhibition has shown dose-dependent anti-proliferative effects in HSC-2 cells and SAS cells xenograft models, with additional inhibition of vascular endothelial growth^30^, suggesting MDK inhibition as a potential approach to disrupt age-associated multicellular communities such as CM07.

Our study includes several limitations. Age in HPV-negative HNSCC is intertwined with other exposures and clinical factors, including smoking and comorbidities, which may contribute to some of the observed differences. Although correlation of the populations in the CM07 community, their marker genes, and specific ligand-receptor pairs were identifiable across independent datasets, functional perturbation studies are needed to establish whether these interactions are causal. It remains unclear whether aging acts primarily through stromal remodeling to reshape malignant cell states, or if age-associated malignant programs actively recruit and sustain a supportive stromal environment, for which perturbation studies would be required to establish mechanism.

Our study provides a scalable computational framework for the interrogation of large single cell RNA-seq datasets and identifies an age-associated community of tumor, stromal, and myeloid cells that support a model in which aging in HPV-negative HNSCC involves coordinated enrichment of invasive epithelial states, supported by matrix remodeling and stromal-associated immunosuppressive myeloid cells. Together, these findings suggest that aging in HPV-negative HNSCC is best understood as a coordinated reorganization of the cellular neighborhoods that support malignant progression.

## Methods

### Integration and clustering

Seven single-cell datasets were downloaded from the GEO database (Peng^31^: GSE172577, Choi^32^: GSE181919, Cillo^33^: GSE139324, Quah^14^: GSE225331, and Kurten^34^: GSE164690, Puram^6^: GSE103322) and preprocessed individually using Seurat^35^ to identify cell types at the compartment level pre-integration. Default library size normalization with a scaling factor of 10,000 and logarithmic transformation was applied. Cells were scored for HPCAD^36^ and blueprint^37^ database cell type signatures using singleR^38^. After individual dataset preprocessing, SCTransform was used to normalize raw counts from each object, and 3000 integration features were used for integration with the ‘RPCA’ method. After integration, the primary tumor cells from each major cell type compartments were reclustered separately to identify 89 high resolution clusters. K2taxonomer^39^ was used to hierarchically cluster the 89 clusters with 400 bootstraps. Marker genes for each cluster were identified using Seurat’s FindAllMarkers with test.use = ‘MAST’, max.cells = 2000, and latent.vars = ‘dataset’. Signatures were identified for each cluster as genes with avg_log2FC > 1 and p_val_adj < 0.05.

### Epithelial cell characterization

Epithelial cells were selected from the nonimmune compartment clustering at 0.2 and subset to only include primary tumor cells. They were reclustered using 10 dimensions at a resolution of 0.4, identifying 18 sub-clusters. Marker genes for each cluster were identified using Seurat’s FindAllMarkers with test.use = ‘MAST’^40^ and latent.vars = ‘dataset’. Signatures were identified for each cluster as genes with avg_log2FC > 1 and p_val_adj < 0.05. Overenrichment analysis was performed on each signature geneset with the KEGG^41^, hallmark^42^, and reactome^43^ pathways, accessed through the msigDBR^44^ package and using hypergeometric testing via the hypeR^45^ package. Spatial signatures were taken from Arora et al.^22^, and Seurat’s AddModuleScore was used to score all epithelial cells with the signatures.

### Differential expression analyses

Differential expression was performed using two methods to ensure robustness. Pseudobulk differential expression was performed for each cellular compartment. The intersection of genes across all datasets was used. For each cellular compartment, we aggregated single-cell counts at the patient level (pseudobulk), summed raw counts per patient using Seurat’s AggregateExpression, and excluded genes if they had zero counts in more than one-third of patients or if their average normalized expression was ≤ 0.5 within the cluster. Counts were normalized across patients using EdgeR’s^46^ scaling factors based on library size. We then applied a quasi-likelihood negative binomial generalized linear model with the design matrix ∼ age + sex + dataset to control for technical batch (dataset) and biological sex while testing for age effects.

We also used a single-cell random effect method to model of gene expression with age. For each cluster, at least one third of patients had to have non-zero counts per gene, at least 5% of cells in the cluster had to express the gene, and the average normalized expression of the gene per cluster had to be >0.1. MAST’s^40^ two-part zlm function was used to fit a zero-inflated generalized linear mixed hurdle model for each gene. For each gene, a conditional regression is used to fit the continuous gene expression values and a discrete model is used to fit the zero vs non-zero values. In both models, age, sex and gene detection number are included as fixed effects, while patient and dataset are included as a nested random effect to account for pseudoreplication bias and dataset batch effects. Options method=’glmer’ was used for model estimation, ebayes=FALSE and fitArgsD=list(nAGQ = 0) were used for a faster Laplace approximation, and strictConvergence = FALSE was used to relax optimizer criteria.

### Cell Type Composition Analysis

The atlas was split into old and young categories with patients <=55 classified as young and patient >55 classified as old. DASeq^47^ was used to identify which cells were enriched in old vs young patients on the PCA embeddings. Multiple neighborhood sizes were tested between 20-1000 by intervals of 20.

### TCGA and CPTAC analyses

The TCGA HNSCC samples were filtered to only include patients that were HPV-negative (n = 421). GSVA^48^ was used to score each patient with the 89 cell type atlas signatures composed of genes with > 1 avg_log2FC and fdr < 0.05 for a given cluster, derived using the Seurat FindAllMarkers function with the MAST model controlling for dataset and using 2000 cells per cluster. Multivariable Cox proportional hazards models were used to assess the association between each signature and overall survival, adjusted for age at index. Multiple testing correction was applied using false discovery rate (FDR) adjustment across all signatures. GSVA scores of the cell type signatures within TCGA were associated with each other using a linear model while controlling for sex. In CPTAC, GSVA was similarly used to score each patient with the 89 cell type atlas genesets. These GSVA scores were correlated to specific protein abundances using Pearson correlation.

### Tan data

The Tan et al.^21^ bulk spatial dataset was downloaded from the GEO database via accession number GSE290057. GSVA^48^ was used to assign atlas cluster signature values to Tan’s bulk transcription data on the tumor and TME samples separately. A linear regression was used to regress the gsva cluster signature score on age while controlling for sex, on the tumor and tme samples separately. To test for atlas cluster signatures association with survival, a survival analysis was performed using disease free survival and the gsva cluster scores on the TME and Tumor samples separately.

### Community detection

CoVarNet^16^ was used to identify cooccurring communities of cells. The 89 cell type clusters were used as cell type annotation. Cells from Cillo et al. were removed from the analysis as they only contained immune cells, and cell types with less than 100 cells were also removed. NMF was applied to the frequency matrix using the nsNMF method with k ranks between 2 and 30. The cophenetic correlation coefficient (CCC) was maximized at k = 20, which we used as the optimal rank.

A partial correlation network was created based on cell type proportions. Atlas cells were filtered to include only primary tumor cells and the Cillo dataset was removed as it contains only immune cells. The 89 high resolution cell types were used as annotation, and only included if the cell type had cells from more than 10 patients. The proportion of cells per cell type per patient were calculated. A hurdle model was used to to regress out dataset-level effects. A logistic model was used to model the zero vs non-zero proportions, and a beta regression was used to model the continuous proportions. For each cell type, we compute the model-expected value E[Y] under the hurdle model as the probability of being nonzero times the conditional nonzero mean. The combined residual is then the observed value minus the expected value.

The RSNet^17,49^ GGM^50^ model was then used to create a network with the resulting residuals with 500 iterations, bootstrapping, and method = ‘B_NW_SL’. A 95% confidence interval was found, and edges with fdr < 0.05 were filtered out for the graph visualization.

### Cell cell communication

CellChat^51^ was used to examine ligand receptor interactions between the 89 clusters on each patient separately, to only identify interaction occurring in cell types present in the same patient, thus eliminating false positives. Genes had to be expressed in at least 1% of the cluster in each patient to be considered, and each cluster had to have at least 10 cells in a patient to be considered. Cellchat’s truncatedMean method was used with trim = 0.2. Custom scripts were used to visualize aggregated predicted signaling across patients in heatmaps and bubble plots that can be accessed through the CellChatCustomPlots package at https://github.com/lkroeh/CellChatCustomPlots.

### Spatial Validation

The Arora et al.^22^ Visium dataset was downloaded from the GEO database via accession number GSE208253. AddModuleScore was used to score each spot for the 89 cell type signatures. Correlation of signatures, or of signatures and genes across samples was calculated using a linear regression while controlling for sample as a fixed effect.

### Mouse Validation

The mouse single cell dataset was downloaded from the GEO database. The dataset was clustered at resolution 0.4, and clusters 5 and 12 were selected as myeloid cells which were then isolated and reclustered at resolution 0.6. AddModuleScore was used to score the myeloid cells with the Mac5_FOLR2 atlas signature. Clusters 14 and 15 were selected as fibroblasts and endothelial cells and reclustered at resolution 0.2. AddModuleScore was used to score the cells for iCAF and mCAF signatures.

### Statistics and Reproducibility

95% confidence intervals were used for all linear regression models. FDR-corrected hurdle p-values were used for all differential expression testing through *MAST*.

Multiple testing correction was applied using false discovery rate (FDR) adjustment for any modeling that included multiple tests such as survival analyses and cell type signature correlations.

Cell-cell communication analyses used empirical p-values in dotplots showing specific ligand-receptor interactions between cell types. P-values are calculated through the *CellChat* package as the fraction of (cell type label) permuted interaction scores that are as large or larger than the original score.

## Acknowledgements

The results published/shown here are in part based upon data generated by the TCGA Research Network: https://www.cancer.gov/tcga.

This work was supported in part by NIH NIDCR award F31DE033292, NIH NIDCR award R01DE033519; NIH NIDCR award R01DE031831, NIH NCATS BU-CTSI award 1UL1TR001430 (SM), Find the Cause Breast Cancer Foundation (SM).

Research reported in this publication was supported by the National Institute Of Dental & Craniofacial Research of the National Institutes of Health under Award Number

F31DE033292. The content is solely the responsibility of the authors and does not necessarily represent the official views of the National Institutes of Health.

## Supplementary Figure Legends

**Supplementary Figure 1:**
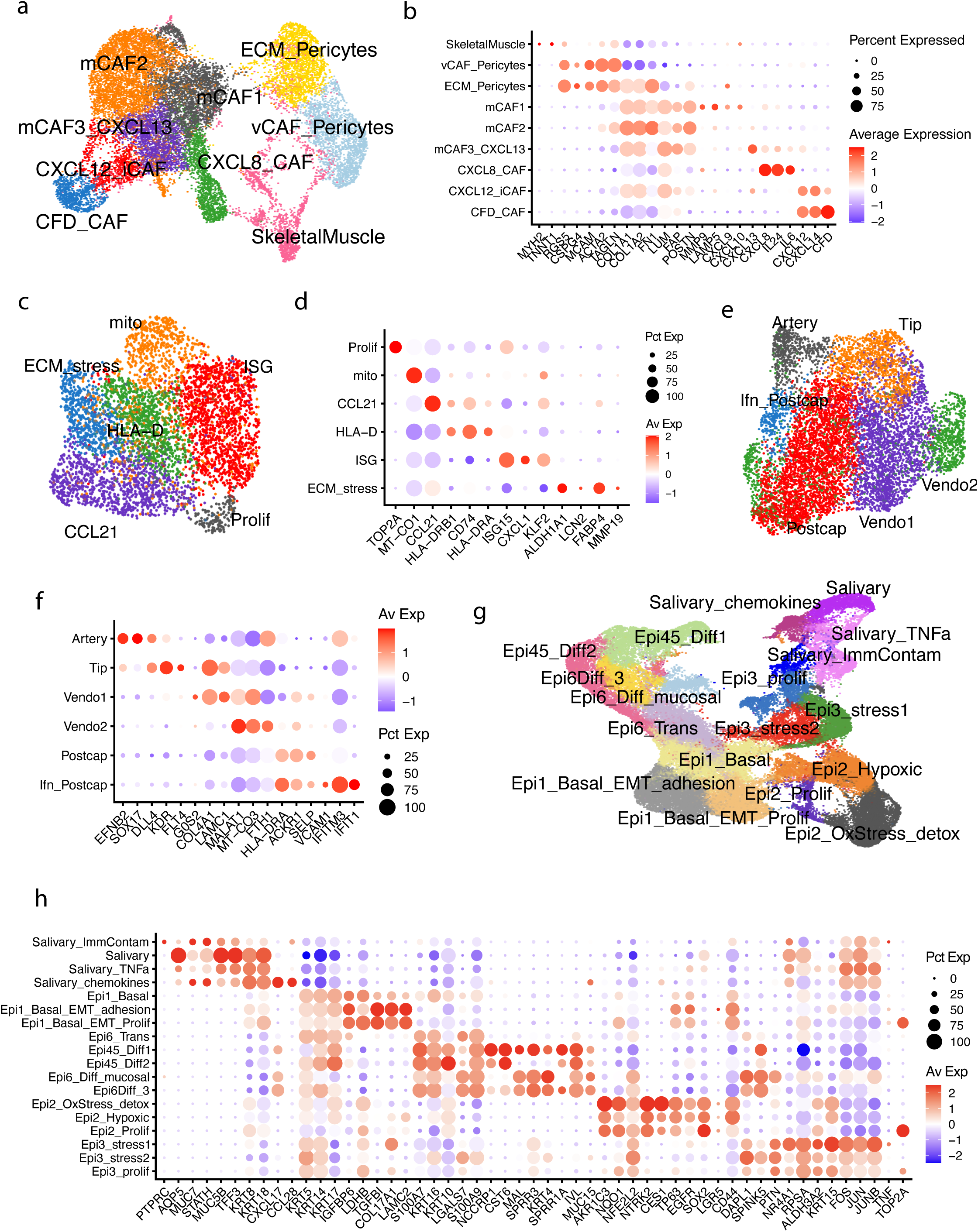
Summary of subclustering of each nonimmune compartment. a) UMAP of mesenchymal cells reclustered at resolution 0.8 for a total of 10 subclusters. b) Dot plot of marker genes for each cluster. c) UMAP of lymphatic endothelial cells reculstered at resolution 0.4 for a total of 6 clusters. d) Dot plot of marker genes for each subcluster. e) UMAP of vascular endothelial cells reclustered at resolution 0.4 for a total of 6 clusters. f) Dot plot of marker genes for each subcluster. g) UMAP of Epithelial and Salivary cells reclustered at resolution 0.4 for a total of 18 subclusters. h) Dot plot of marker genes for each subcluster. For all dot plots, color represents scaled average normalized gene expression, and dot size represents percent of cells per cluster expressing the given gene.

**Supplementary Figure 2:**
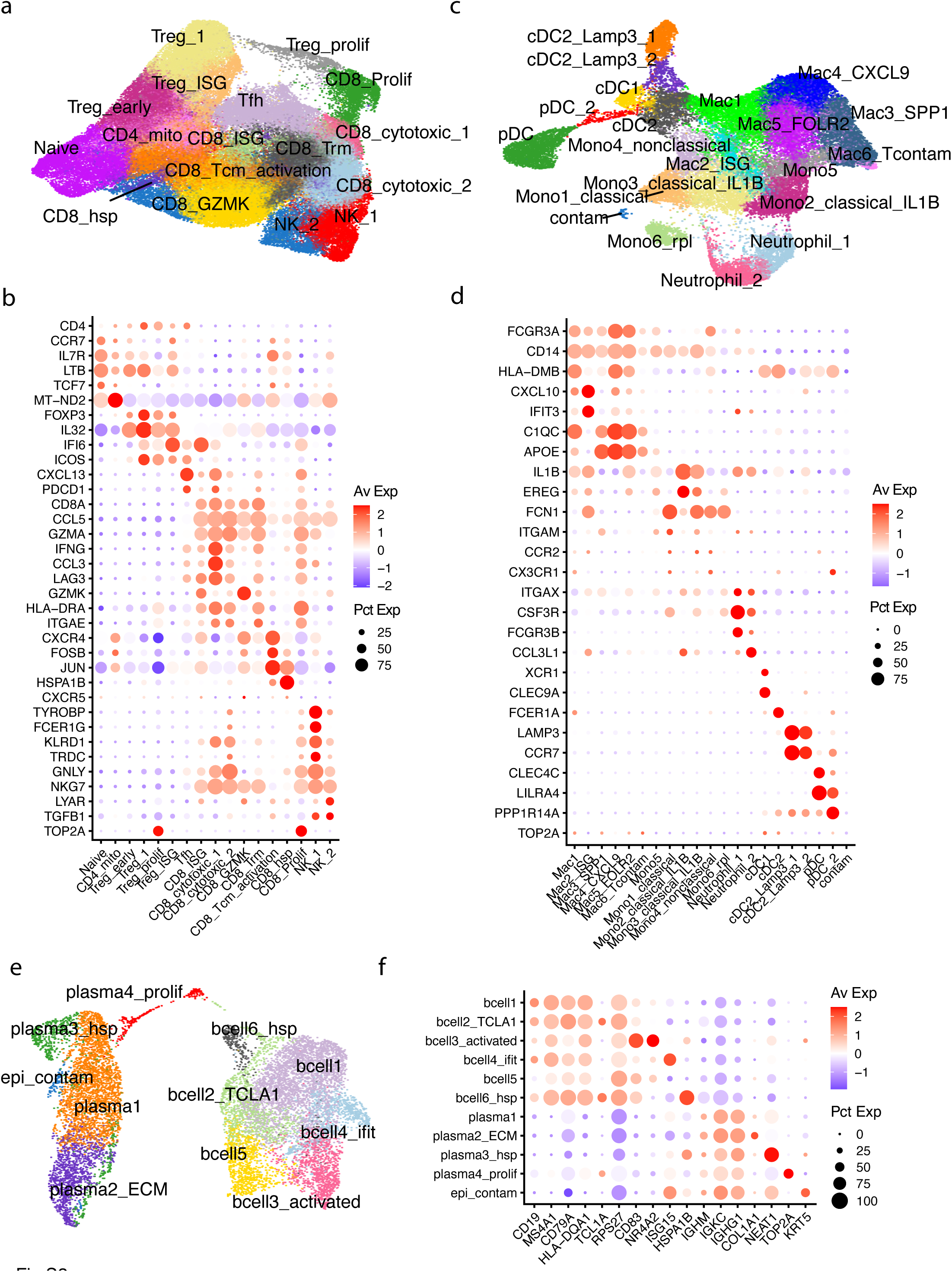
Summary of subclustering of each immune compartment. a) UMAP of T and NK cells reclustered at resolution 0.6 for a total of 17 subclusters. b) Dot plot of marker genes for each subcluster. c) UMAP of myeloid cells reclustered at resolution 0.8 for a total of 21 subclusters. d) Dot plot showing marker genes for each subcluster. e) UMAP of B and plasma cells reclustered at resolution 0.4 for total of 11 subclusters. f) Dot plot of marker genes for each subcluster. For all dot plots, color represents scaled normalized gene expression, and dot size represents percent of cells per cluster expressing the given gene.

**Supplementary Figure 3.**
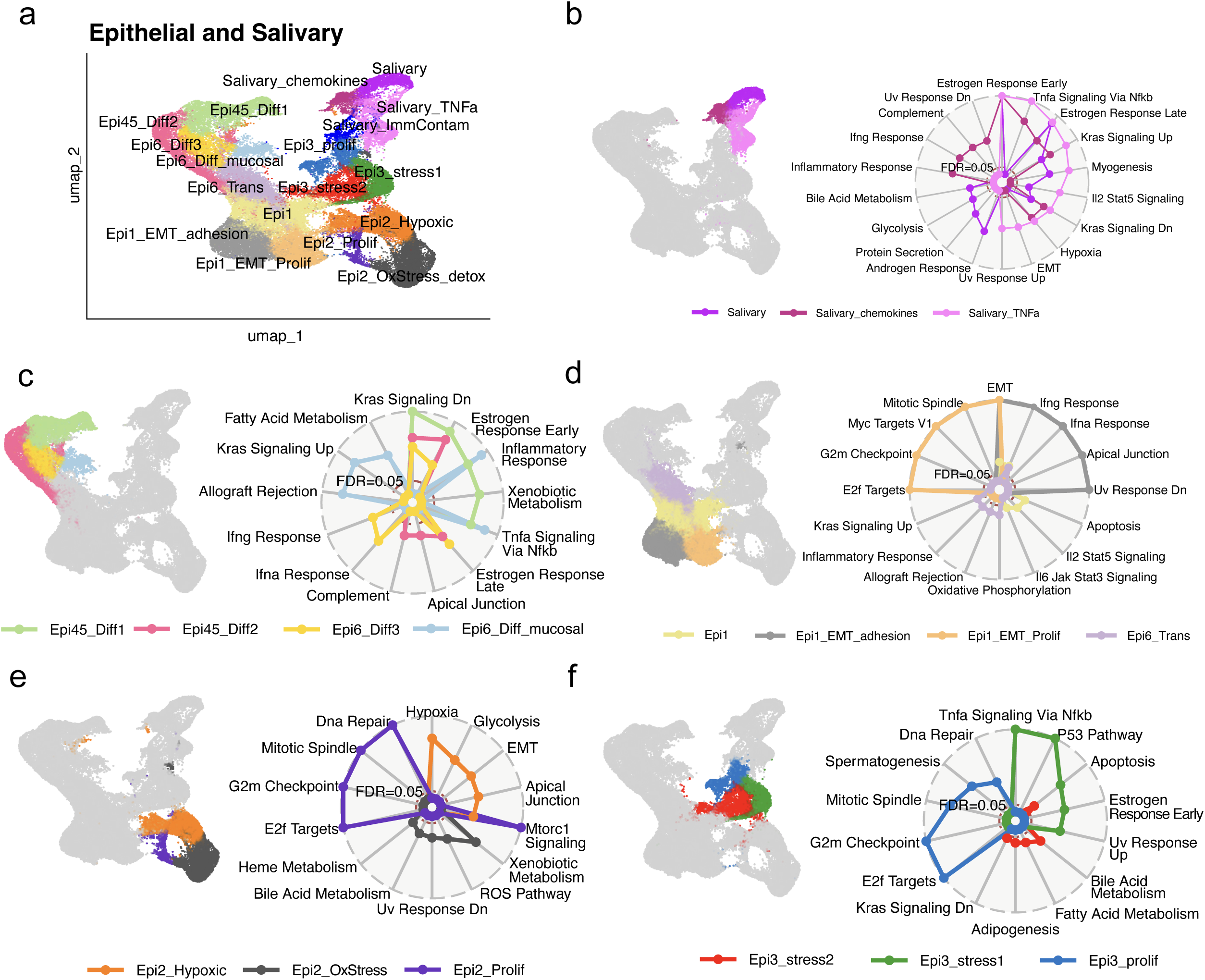
Epithelial compartment characterization. a) UMAP of the epithelial and salivary compartment reclustered at resolution 0.4 for a total of 17 cell sub-types. b-f) UMAP with Salivary, Differentiated, Basal-like squamous, Epi2, and Epi3 sub-clusters highlighted with spider plots showing overrepresentation analyses via hypergeometric test in the hallmark dataset (right). The top 10 overrespresented pathways with FDR < 0.05 for each group are shown.

**Supplementary Figure 4.**
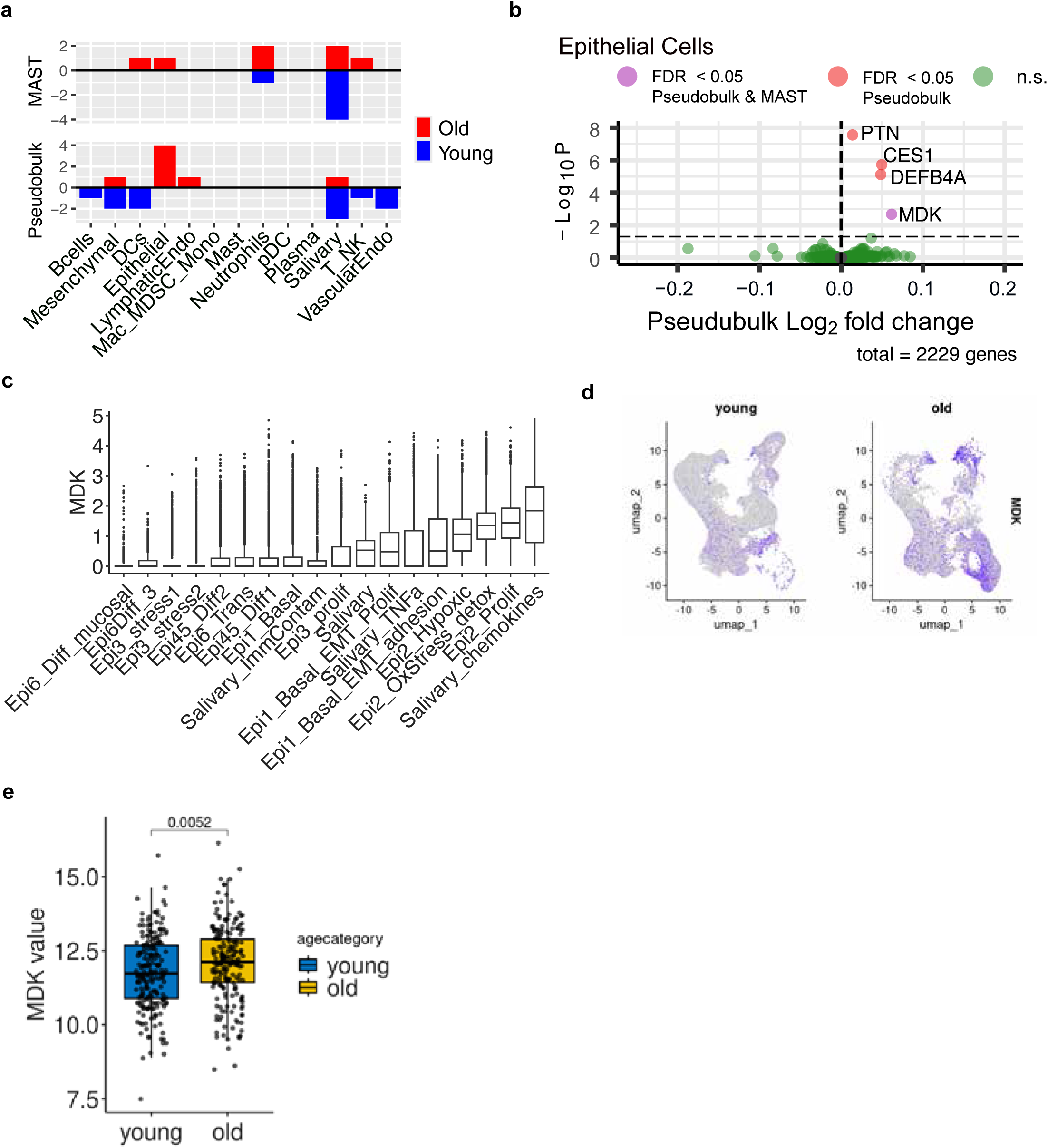
Differential expression analysis. a) Barplots showing number of significantly differential genes (fdr < 0.05) in old (red) or young (blue) patients across cell types using MAST (top) or pseudobulk (bottom) methods. b) Volcano plot showing pseudobulk differential expression results within the epithelial cells. Red dots represent significantly changing genes with fdr < 0.05. The pink dot shows the single gene also significantly (fdr < 0.05) differential in the MAST analysis, MDK. c) Boxplots showing normalized MDK expression across the epithelial and and salivary clusters. d) UMAP showing MDK expression in young (< 65 years) and old (>= 65 years) patients. e) Boxplot showing normalized MDK expression in young vs old patients from the HPV-negative TCGA dataset. Wilcxon test, pval = 0.0052.

**Supplementary Figure 5:**
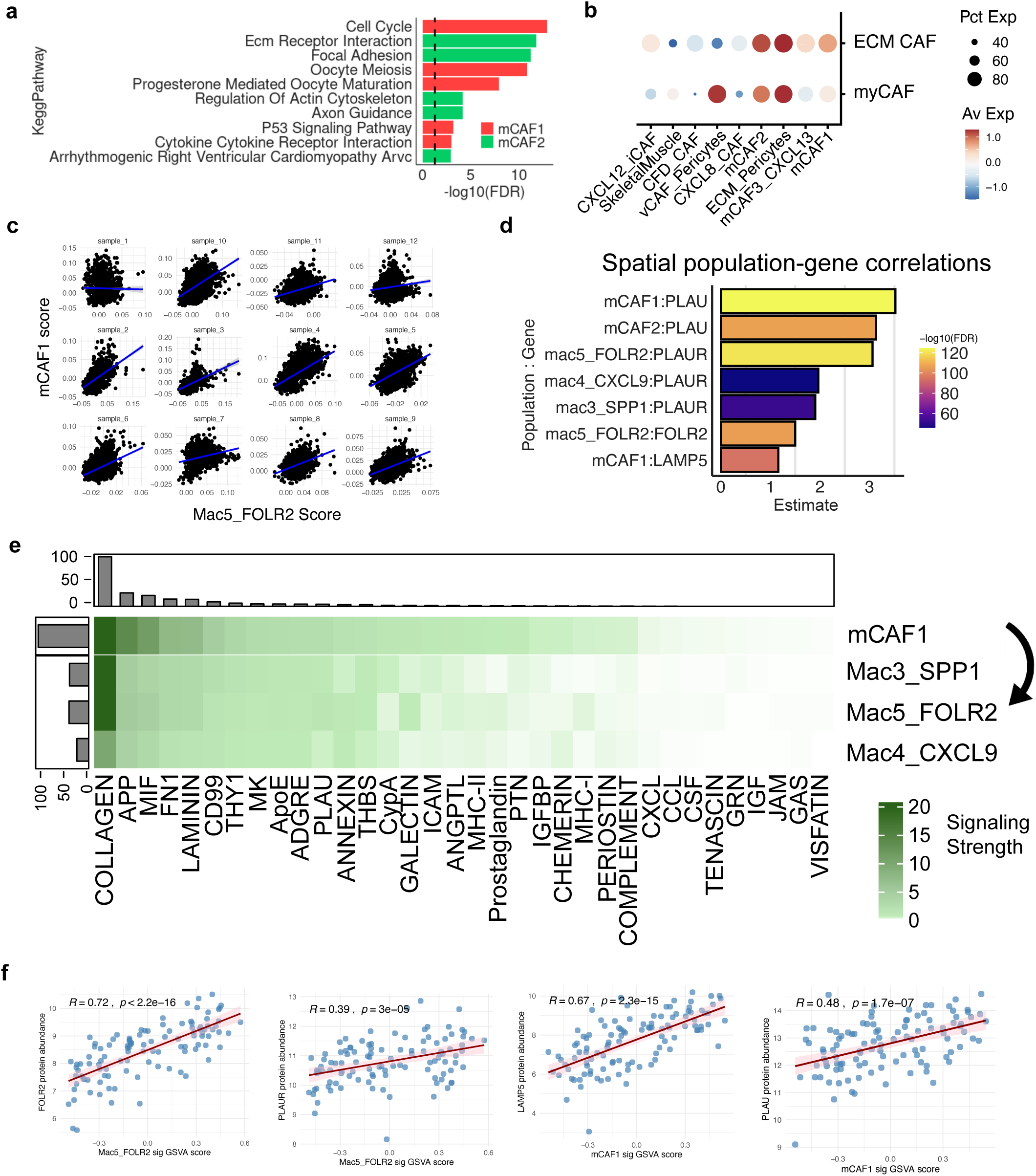
CM07 population supporting information. a) Overrepresentation analysis of the mCAF1 (red) and mCAF2 (green) clusters via hypergeometric test in the Kegg pathway dataset. b) Dotplot showing myCAF and ECM CAF signature enrichment across mesenchymal clusters. c) Correlation of the mCAF1 and Mac5_FOLR2 signatures in the 12 Arora spatial patients. d) Estimates for celltype:gene correlations (y-axis) in the spatial dataset. Bars are colored by fdr. e) Heatmap showing signaling pathways sent from mCAF1 to Mac3_SPP1, Mac4_CXCL9, and Mac5_FOLR2 populations. f) Pearson correlation of protein abuandances (y-axis) of population marker genes and corresponding atlas signature GSVA scores from CPTAC HNSCC cohort.

